# Revealing the chassis-effect on a broad-host-range genetic switch and its concordance with interspecies bacterial physiologies

**DOI:** 10.1101/2023.02.27.529268

**Authors:** Dennis Tin Chat Chan, Geoff S. Baldwin, Hans C. Bernstein

## Abstract

Broad-host-range synthetic biology is an emerging frontier that aims to expand our current engineerable domain of microbial hosts for biodesign applications. As more novel species are brought to “model status”, synthetic biologists are discovering that identically engineered genetic circuits can exhibit different performances depending on the organism it operates within, an observation referred to as the “chassis-effect”. It remains a major challenge to uncover which genome encoded and physiological biological determinants will underpin chassis effects that govern the performance of engineered genetic devices. In this study, we compared model and novel bacterial hosts to ask whether phylogenomic relatedness or similarity in host physiology is a better predictor of toggle switch performance. This was accomplished using comparative framework based on multivariate statistical approaches to systematically demonstrate the chassis-effect and characterize the performance dynamics of a genetic toggle switch operating within six Gammaproteobacteria. Our results solidify the notion that genetic devices are significantly impacted by host-context. Furthermore, we formally determined that hosts exhibiting more similar metrics of growth and molecular physiology also exhibit more similar toggle switch performance, indicating that specific bacterial physiology underpins measurable chassis effects. The result of this study contributes to the field of broad-host-range synthetic biology by lending increased predictive power to the implementation of genetic devices in less-established microbial hosts.

## INTRODUCTION

While the collection of modular genetic parts (e.g., promoters, reporter proteins, etc.) has grown rapidly over the years, the number of genetically tractable “domesticated” microbial hosts, or chassis, has remained relatively small. Indeed, despite a populous collection of cultivable bacteria readily available in culture collections^1^, contemporary biodesign efforts still preferentially employ the same handful of model organisms (e.g., *Escherichia coli*, *Saccharomyces cerevisiae*, *Bacillus subtilis* and *Pseudomonas putida*). While these model species are easy to work with, they do not always serve as the most optimal chassis for the intended objectives and could instead limit the potential of synthetic biology applications^2^. Broad-host-range (BHR) synthetic biology is an emerging field seeking to expand our engineerable domain beyond that of traditional model organisms, and in doing so, allows us to take advantage of the rich phenotypic diversity of naturally evolved microorganisms^3–5^ to construct more sophisticated bespoke systems.

The number of studies promoting novel microbes for synthetic biology applications is increasing^6–12^, demonstrating that the field of BHR synthetic biology has gained considerable traction. As synthetic biologists continue to explore the chassis-design space, we (re-)discover that genetic circuits do not always maintain similar functional fidelity across hosts. Indeed, previous studies have shown that the same genetic circuit can exhibit significantly different behavior depending on the host environment it is operating within, an observation termed the “chassis-effect”^13–16^. The chassis-effect may hinder the accurate prediction of function from genetic composition alone^17, 18^, which can be disarming and lead to additional costly repetitions of the design-build-test cycle. The chassis-effect can also render any optimizations of a circuit done in the context of a “design” host (typically a cloning-optimized strain) obsolete once transformed into the cellular environment of the destination host. This often discourages the use of non-model organisms. On the other hand, previous literature has demonstrated how the chassis-effect can be exploited to expand the functionality and properties of circuits^15, 16^. In this perspective, the host is viewed as a part that can be used to tune circuit function. The chassis-effect can therefore act as an obstacle, but also as an opportunity. However, a predictive understanding of which specific biological properties underpin observable chassis-effects is lacking, representing a major knowledge gap that has been left unanswered due to the defaulting to the use of model organisms. Filling this knowledge gap will not only help mitigate the degree of uncertainty caused by the chassis effect, but also provide more predictive power to BHR synthetic biology applications and contribute to broadening the design space available for biodesign applications.

Studies in the field of biosynthetic gene cluster expression^19^ and microbial community engineering^20^ have shown that similarities in phylogeny and genotypic profiles can accurately predict metabolic phenotype, which suggests genome relatedness could be a potential predictor of genetic circuit performance. On the other hand, the functional phenotype of expression plasmids and genetic devices has been shown to be coupled to physiological metrics such as growth rate^21, 22^, gene copy number^23, 24^, codon usage bias^25, 26^ and growth burden^27, 28^ in a number of studies. These studies have however, only considered a single or select combination of physiology metrics as explanatory variables of phenotype within a single model chassis. Here, we detail a comprehensive study that takes account a multitude of explanatory variables within a comparative framework that includes model and non-model bacterial chassis. We systematically demonstrate the chassis-effect by characterizing the performance dynamics of a genetic toggle switch, as an example of an engineered gene circuit, within six different Gammaproteobacteria species. As our major guiding research question, we ask whether phylogenomic relatedness or similarity in assorted host physiology metrics is a better predictor of toggle switch performance. Due to the more extensive documentation on the coupling of host physiology and gene expression, we hypothesized that variations in the physiology between hosts will more robustly predict variations in performance of a genetic device between hosts, compared to phylogeny. This hypothesis was tested in our six bacterial hosts operating the same genetic toggle switch under identical growth conditions, using a combinatorial multivariate statistical approach.

## RESULTS

### Comparative physiology and phylogeny between hosts

Measuring biological determinants of the chassis-effect requires a standardizable, BHR device that can be ported across multiple species. We therefore built a tractable experimental platform by transforming six Gammaproteobacteria species with a genetic toggle switch built onto a pSEVA231-derived (Kanamycin selection marker and pBBR1 origin of replication) vector (Fig. 1a, 1b) within the Biopart Assembly Standard for Idempotent Cloning (BASIC) cloning environment^29, 30^, yielding plasmid pS4. The toggle switch consists of two inducible operons acting as antagonistic expression cassettes reported by either mKate or sfGFP fluorescence signals. This mutual inhibitory motif allows cells carrying the toggle switch to achieve two distinct ON states in the presence of either anhydrotetracycline (aTc) or L-Arabinose (Ara) input inducers. The toggle switch was successfully transformed into the following six species: *Escherichia coli* DH5α (*E. coli*), *Halopseudomonas aestusnigri* VGOX14 (*H. aestusnigri*), *Halopseudomonas oceani* KX20 (*H. oceani*), *Pseudomonas deceptionensis* M1 (*P. deceptionensis*), *Pseudomonas fluorescens* SBW25 (*P. fluorescens*) and *Pseudomonas putida* KT2440 (*P. putida*). Members of the *Pseudomonas* and *Halopseudomonas* genus were specifically chosen because of their known robustness and metabolic capabilities^31, 32^, making them attractive candidates for biotechnology applications and for comparing bacterial ecophysiology.

**Figure 1.**
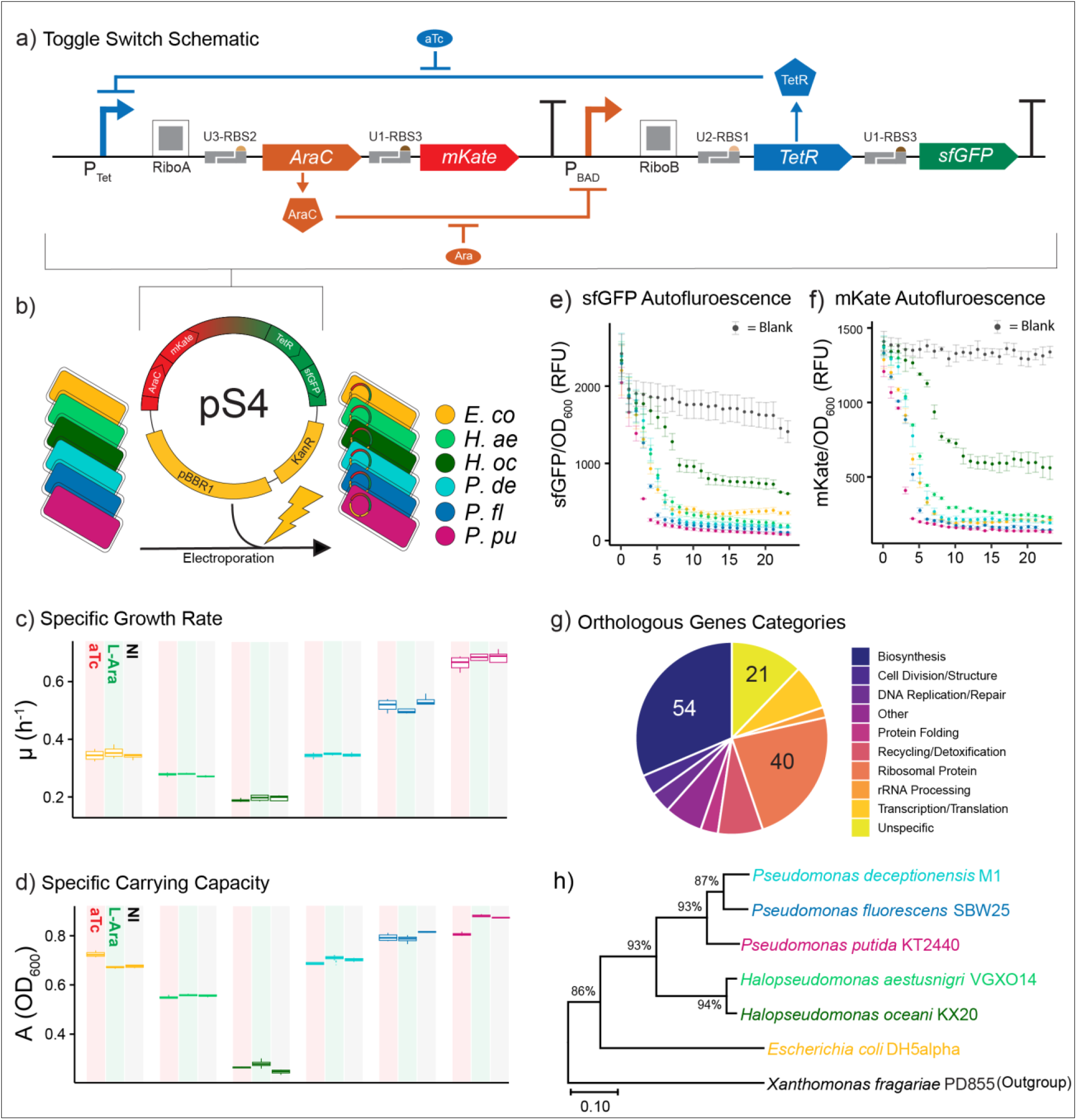
The broad-host range toggle switch pS4 was introduced into six distinct hosts from the Gammaproteobacteria class. a) Schematic of Ara and aTc inducible toggle switch. In presence of aTc, upstream cassette is upregulated, leading to production of mKate reporter protein and AraC repressor, the latter binds to its cognate PBAD promoter to downregulate the downstream cassette. In presence of Ara, sfGFP and TetR repressor is expressed, leading to downregulation of the upstream cassette. RiboA and RiboB are autocatalytic ribozyme insulators that, once transcribed, cleave the upstream 5’ end of the expressed mRNA, standardizing the length of the UTR-5’ region in transcribed mRNA molecules. U*n*-RBS*n* are BASIC linkers with a ribosome binding site (RBS) in their adapter region. Number in U*n* indicates the BASIC linker family (1, 2 or 3) while the number in RBS*n* indicates relative translational strength of the RBS, from 1 (weakest) to 3 (strongest). b) The toggle switch was assembled into a BASIC integrated pSEVA231 vector, with a kanamycin selection marker and pBBR1 origin of replication, resulting in plasmid pS4. Plasmid pS4 was transformed by electroporation into six bacterial Gammaproteobacteria hosts. Hosts are color coded; all subsequent panels follow the same color code. *E. co* = *E. coli*, *H. ae* = *H. aestusnigri*, *H. oc* = *H. oceani*, *P. de* = *P. deceptionensis*, *P. fl* = *P. fluorescens*, *P. pu* = *P. putida*. c) Specific growth rate (µ) and d) carrying capacity (A) of WT strains in presence of aTc (10 ng/mL, red columns) and Ara (20 mM, green columns) and no inducer (NI, grey columns). Error bars show standard error of the mean (n = 3 biological replicates, with 4 technical replicates each). sfGFP e) and mKate f) autofluorescence normalized by OD600 over time by WT strains in absence of inducer. Blank is autofluorescence of wells containing only LB media. g) The 172 single-copy genes in the Gammaproteobacteria Hidden Markov Model set from GToTree used for phylogeny inference grouped by functional annotation. Number of genes in the largest grouped are indicated. h) Phylogenomic tree inferred from multi-locus sequence alignment of concatenated gene hits. *Xanthomonas fragariae* PD855 was chosen as outgroup. Scale bar indicates the number of amino acid substitutions per site between two sequences. Tree was inferred using Neighbor-Joining method in MEGAX. The percentage of replicate trees in which the associated taxa clustered together in bootstrap test (1000 replicates) are shown next to the branches.

A preliminary growth characterization of the wild-type (WT) strains in absence and presence of Ara and aTc inducers was performed (Fig. 1c, 1d). Each species revealed distinct growth physiologies, allowing for quantitative comparison of the relative contribution that a host’s physiology plays on device performance. In absence of any inducer, *P. putida* achieved the highest maximum specific growth rate (µ = 0.67 h^-1^ ± 0.01). The other two *Pseudomonas* spp. exhibited lower growth rates compared to *P. putida* but reached similarly high specific carrying capacities (A, Fig. 1c, 1d). In contrast, the two *Halopseudomonas* members showed approximately 2 to 3-fold lower µ values as compared to *P. putida*. Given the documented impact of growth rate on functional output of genetic devices, such as how rapid cell division results in dilution of gene expression machinery and expression products^21^, the range of growth dynamics observed lead us to predict that a measurable chassis-effect would be observed among our hosts. No appreciable difference in growth rate or carrying capacity was observed when the WT strains were grown in presence of either inducer, suggesting negligible levels of toxicity. Furthermore, none of the species had significant levels of autofluorescence (Fig. 1e, 1f).

The relative performance of the genetic toggle switch is expected to be in part related to the genomic potential of the host. For example, genes encoding for the specific transcriptional and translational proteins used to express the toggle switch’s machinery will ultimately underpin performance. We used a phylogenomic approach to ask if hosts that share more genome relatedness will also share similar device performances, employing the GToTree program^33^ due to its ease of use and reproducibility. The Gammaproteobacteria Hidden Markov Model (HMM) set containing 172 orthologous single-copy genes fit for phylogeny inference in Gammaproteobacteria from the GToTree was used to infer phylogeny between our hosts (Fig. 1g). The set included genes encoding for ribosomal proteins, chaperones and transcription and translation proteins, which together are partly responsible for shaping the gene expression landscape of each host such as steady-state concentration of transcriptional/translational machinery and turnover rate. Hosts with higher sequence similarity in these specific genes could therefore be expected to have similar device performance. Genomes with the lowest gene hits identified after filtering for redundancy and length belonged to *Xanthomonas fragariae* PD855 (outgroup) and *P. fluorescens*, both with 159 gene hits, while the other five genomes retained between 166 to 170 genes (Supplementary Table S1). A phylogenomic tree clustered the hosts into distinct clades according to their genus, with the *Halopseudomonas* clade being more closely related to the *Pseudomonas* clade and *E. coli* being distant from both (Fig. 1h). The growth characterization demonstrated that closely related hosts also shared similar growth physiologies. These results set the stage to comparatively investigate how the performance of the genetic toggle switch relates the relative contribution of species-specific physiology as compared to phylogenomic relatedness – (i.e., more similarly evolved genomes) – and to ask whether host physiology or phylogeny is a more robust predictor of differences in the performance of a genetic toggle switch between hosts.

### Quantifying the chassis-effect

The toggle switch was functional across all six hosts. Induction kinetic and expression dynamic assays were performed to quantify toggle switch performance operating under the cellular context of each chassis. Significant differences in performance metrics were observed between chassis, establishing a clear and quantifiable chassis-effect (Fig. 2).

**Figure 2.**
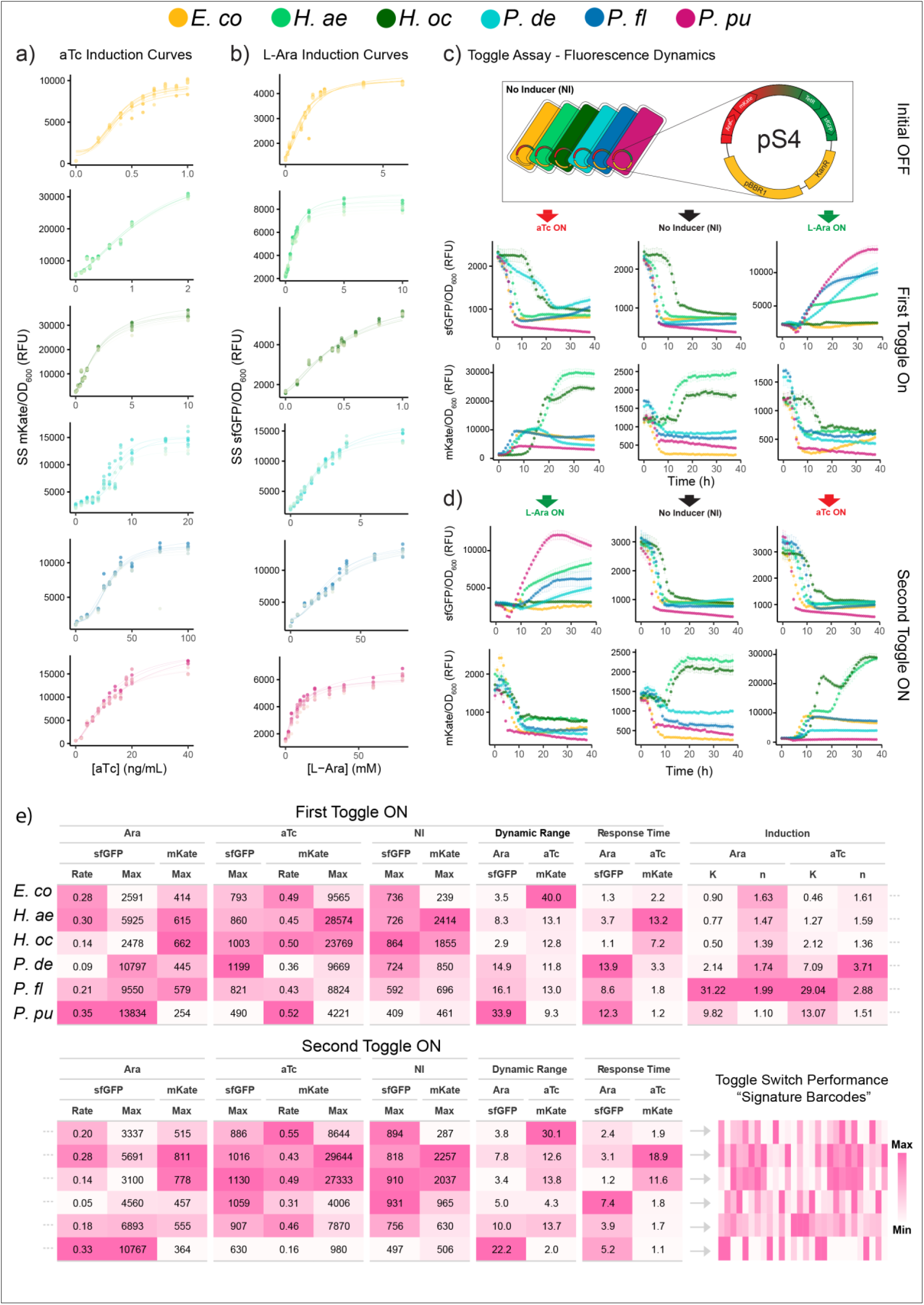
The chassis-effect as observed through heterogeneity of performance metrics of the ON/ON genetic toggle switch pS4. a) aTc and b) Ara induction curves of pS4-carrying hosts with late phase steady-state (SS) fluorescence intensity (mKate and sfGFP respectively) normalized by OD600 as response variable on the y-axis. Fitted Hill function are shown. Color shades represent technical replicates (n = 8). Hosts are color coded according to legend on figure top. c) Normalized sfGFP and mKate fluorescence dynamics over time of pS4-carrying hosts toggled from initial OFF state (no inducer) to ON state (First Toggle ON) with 20 mM Ara (right, green arrow) and 10 ng/mL aTc (left, red arrow). No inducer (NI) control was also included (middle, black arrow). d) Cells were toggled by washing away inducer twice and diluted to media with opposite inducer at same concentration (Second Toggle ON). e) Summary of performance metrics estimated from induction and toggle assay experiments for each host, with each column representing a specific performance metric. The color scale is relative to each column, allowing a visual comparison of host similarity within a metric and the abstraction of a unique “signature barcode” that profiles comparative toggle switch performance of each chassis. Grey dots denote continuation of table. The SS fluorescence intensity representing leakage was quantified by averaging normalized fluorescence over a time window of at least 20 hours at late phase. Rate = maximum rate in unit of h^-1^, SS = steady-state fluorescence at late phase in units of relative fluorescence units (RFU), NI = No Inducer, DR = Dynamic range, Response Time in units of hours, K = activation constant, n = Hill coefficient. Color scale denoting “Max” and “Min” value is relative for each column.

Induction kinetics of the toggle switch was resolved by fitting the Hill equation to Ara and aTc induction response curves to estimate the species-specific activation constants (K_Ara_ and K_aTc_) and Hill coefficients (n_Ara_ and n_aTc_) (Fig. 2a, b). *P. fluorescens* exhibited the largest inducible concentration range for both inducers (K_Ara_ = 31.2 mM ± 2.1 and K_aTc_ = 29.0 ng/mL ± 1.1). *E. coli* and both *Halopseudomonas* spp. revealed some of the lowest respective activation coefficients. The differences in activation constants between and within chassis represents how each host uniquely mediates the concentration and/or binding affinities of free intracellular inducer that is available to allosterically bind its cognate repressor. Furthermore, both sigmoidal responses and damped “titratable responses” between hosts across induction states was observed, but all hosts had a Hill coefficient close to or higher than 1. The Hill coefficients n_Ara_ and n_aTc_ did not differ drastically within hosts, the only exception being *P. deceptionensis*, which exhibited a relatively more step-like response behavior to aTc (n_aTc_ = 3.71 ± 0.53) compared to Ara induction (n_Ara_ = 1.74 ± 0.17). Clearly, the choice of chassis can greatly impact the operational properties of a genetic device, demonstrating the importance of considering the chassis during design stages.

To further quantify the chassis-effect, fluorescence dynamics of each host grown under the same induction state was characterized through a “toggling assay” (Fig. 2c, First Toggle ON). Cells in initial OFF state (no inducer, NI) was diluted to media containing Ara or aTc to toggle each respective promoter ON and cultured for 40 hours before being washed and toggled to the opposite state. To quantify host-specific fluorescence dynamics, maximum rate (Rate) and steady-state (SS, plateau of curves at late phase) fluorescence level was estimated from normalized sfGFP and mKate curves across induction states. We also determined two additional metrics to characterize species-specific toggle switch performance, “dynamic range” (DR) and “response time”, which we define here as the ratio between induced SS fluorescence and non-induced SS fluorescence (DR_sfGFP_ = SS_Ara-sfGFP_/SS_NI-sfGFP_ and DR_mKate_ = SS_aTc-mKate_/SS_NI-mKate_) and the time it takes to reach half of the SS fluorescence (subtracting for time during lag phase), respectively.

Under Ara induction, the three *Pseudomonas* spp. display the highest SS_sfGFP_ values, which reflects the attained SS concentration of sfGFP in the population. *P. putida* attained the highest value (SS_sfGFP_ = 13833 RFU ± 937) and the highest dynamic range (DR_sfGFP_ = 33.85 ± 1.37) (Fig. 2c, 2e). However, the high sfGFP output level from these three chassis was also associated with relatively long response times. *E. coli*, *H. aestusnigri* and *H. oceani* had lower DR_sfGFP_ values, but also shorter response times. Under aTc induction, the highest SS_aTc-mKate_ values was reached by the two *Halopseudomonas* spp., again associated with long response times, although the highest DR_mKate_ value belonged to *E. coli*, with an induced mKate output level 40-fold that of the output level in absence of inducer. Even in the presence of repressor or in absence of activator, some basal level of expression is often observed from an inducible promoter, referred to as leakage^34^. We identify two types of leakages in our toggle switch system: output in absence of any inducer (Fig. 2c, middle) and in the presence of the fluorescent protein’s antagonistic inducer (i.e., sfGFP in presence of aTc and mKate in presence of Ara, Fig 2c, top-left and bottom-right). Under these inducer conditions, the SS fluorescence at late phase was used to quantify leakage, determined by averaging SS fluorescence over a time window of at least 20 hours. Overall, NI control cultures revealed a low amount of basal leakage, indicating similarly tight regulation among hosts. The exception was *H. aestusnigri* and *H. oceani*, which showed mKate SS fluorescence values about 10 and 7-fold higher than *E. coli*, which had the lowest level of mKate leakage. This result suggests that the P_Tet_ promoter is leakier when operating in the cellular context of the two *Halopseudomonas* chassis. We observed up to 4-fold lower mKate leakage in the two *Halopseudomonas* spp. under Ara induction compared to the leakage in absence of inducer, which we attribute to repression of the P_Tet_ promoter by expressed TetR. The wide range of metrics observed from different species cultivated and induced under the same conditions further reinforces the presence of the chassis-effect affecting toggle switch performance.

A system exhibits hysteresis if the state of the system is dependent on its past state(s)^21^. In the context of our toggle switch, hysteresis is observed if the fluorescence dynamics of toggle switch carrying cells (induced at the same concentration) differ because of the cells past induction states. To investigate whether this type of hysteresis is host-specific, cells toggled to Ara/aTc-ON from initial OFF were washed and diluted 200-fold to the opposite induction state (Fig. 2d, Second Toggle ON). When toggled from aTc-ON to Ara-ON, the three *Pseudomonas* spp. consistently experienced a decrease in DR_sfGFP_ values, ranging from 0.78 to 0.42-fold change (Fig. 2e) compared to cells toggled to Ara-ON from an initial OFF state, although this was accompanied with a 0.54 – 0.43-fold change in response time as well. As cells were washed of inducer and diluted, this decrease in expression can be attributed to the cells past induction state, indicating hysteresis. Furthermore, when toggled from Ara-ON to aTc-ON, *P. putida* and *P. deceptionensis* showed a notable 0.37 and 0.21-fold change decrease in DR_mKate_, respectively. *H. aestusnigri* and *H. oceani* showed relatively little differences in fluorescence dynamics despite their past induction state, with most metrics having fold-changes close to 1. These results indicate that, under our induction scheme, some of our chosen hosts are more sensitive to past states of induction than others, in turn demonstrating that the magnitude of hysteresis-effect is host specific.

Summarizing the quantified performance metrics in table format with a color scale allows the set of metrics for each host to be abstracted as a “signature barcode” that serve as an identifier of the hosts performance profile. There is a variable degree of, but clear heterogeneity in the performance metrics among our closely related hosts, despite operating the same genetic device. We conclude that functional parameters of toggle switch activity are highly dependent on the host-context it is operating within, showing that the basis for the chassis-effect stems from unique host-specific properties. We therefore proceeded to characterize the hosts in terms of relevant physiology metrics to encapsulate the unique host conditions each toggle switch is operating under.

### Physiological diversity among hosts

We quantified host-specific parameters known to affect gene expression for each host, which include plasmid copy number (PCN), maximum specific growth rate, carrying capacity, growth burden as a result of propagating and expressing the pS4 plasmid, and finally the codon adaption index^26, 35^ (CAI) for the four coding sequences within the toggle switch. We uncovered that our hosts also display a wide range of physiology metrics, suggesting that physiology could explain, and therein possibly be used to predict, differences in toggle switch performance between bacterial hosts (Fig. 3).

**Figure 3.**
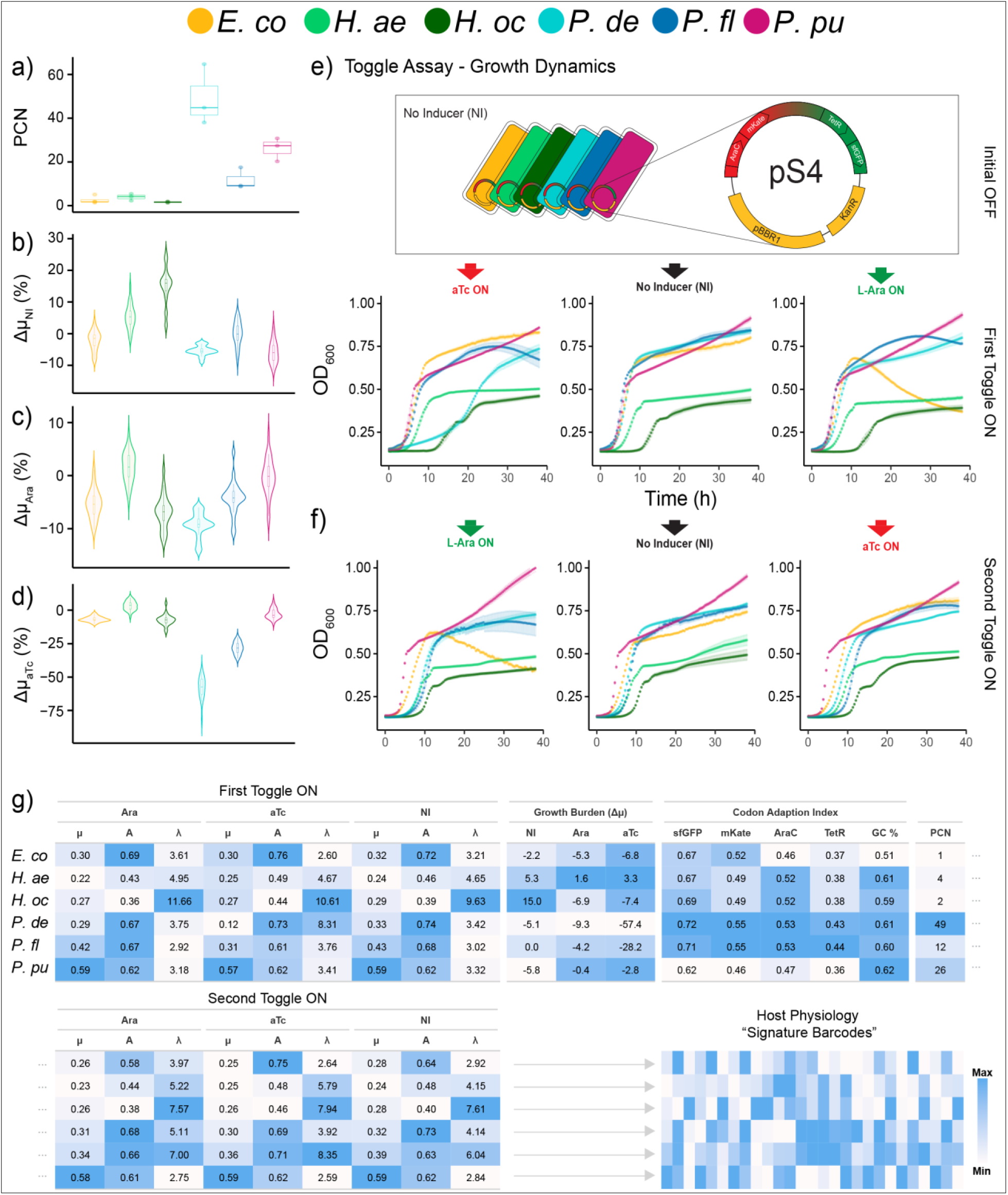
The hosts display a wide range between values among the metrics chosen to capsulate physiology. a) Boxplots of estimated plasmid copy number (PCN) per chromosome of pS4-carrying strains (n = 3 biological replicates each with 4 technical replicates). Hosts are color coded according to legend at top of figure. Violin plots showing b) growth burden (ΔµNI) in absence of inducer, defined as the percent difference in specific growth rate of pS4 carrying strains compared to WT controls. A negative ΔµNI value indicates the pS4 harboring strain has a lower growth rate compared to its WT counterpart. c) Growth burden of Ara induction (ΔµAra) and d) aTc induction (ΔµaTc), defined as the difference in growth rate of Ara and aTc induced pS4-carrying strains compared to the growth rate under NI conditions, respectively (n = 3 biological replicates each with 4 technical replicates). e) Growth dynamics of hosts from toggling assay experiment toggled from initial OFF (no inducer) to ON state (First Toggle ON) and then f) to opposite ON state (Second Toggle ON). g) Summary of quantified physiology metrics for each host, with each column representing a specific metric. The color scale is relative to each column, allowing a visual comparison of host similarity within a metric and the abstraction of a unique “signature barcode” that profiles comparative host physiology of each chassis. Grey dots denote continuation of table. µ = maximum specific growth rate (h^-1^), A = specific carrying capacity (OD600), NI = no inducer, PCN in units of plasmid(s) per chromosome. GC % = total genomic GC content. Color scale denoting “Max” and “Min” values are relative for each column.

PCN, in units of plasmid(s)/chromosome, was determined through a real-time PCR based method^36^. Despite harboring the same origin of replication, a wide range of PCN values was observed among our hosts (Fig. 3a). This shows that PCN is also subject to the chassis-effect, which is in agreement with previous studies characterizing PCN of the same plasmid across microbes^37, 38^. Maintenance of a higher number of plasmids has been shown to cause growth inhibition^39^, which can be exacerbated upon circuit induction^23^, a scenario applicable to the operation of our toggle switch. We therefore quantified the growth burden of maintaining the toggle switch in absence of inducer, given by the Δµ_NI_ metric, which we define as the percent difference in specific growth rate of pS4-carrying strains compared to WT controls. A negative Δµ_NI_ value indicates the pS4-carrying strain has a lower growth rate compared to its WT counterpart. We also define Δµ_Ara_ and Δµ_aTc_ as the percent difference in growth rate of Ara and aTc induced pS4-carrying strains compared to the growth rate under NI conditions, respectively. The *Pseudomonas* spp. maintained the highest number of plasmid copies, with *P. deceptionensis* peaking at a PCN value of 49 ± 12. In absence of inducer, *P. deceptionensis* showed slight but consistent reduced growth compared to their WT counterparts (Δµ_NI_ = -5.4 ± 0.9%) (Fig. 3b). Meanwhile, a surprising result was the consistent increase in growth rate by both *Halopseudomonas* spp., with *H. oceani* pS4 having a specific growth rate 15.0 % ± 3.4 higher than its WT counterpart despite maintaining an additional plasmid, albeit at a relatively low copy number (PCN = 2 ± 1). Upon induction, *P. deceptionensis* experienced the strongest exacerbation of growth burden, signified by the most negative Δµ_Ara_ and Δµ_aTc_ values of -9.2 % ± 0.3 and -58.0 % ± 5.0, respectively (Fig 3c, 3d). *P. fluorescens* was particularly sensitive to aTc induction, with a Δµ_aTc_ value of -28.2 % ± 3.5. The high growth burden of *P. deceptionensis* and *P. fluorescens* is likely attributable to their high PCN. However, *P. putida*, which had the second highest PCN of 26 ± 5, deviated from this trend with Δµ values close to 0. None of the WT strains experienced appreciable growth inhibition when grown in presence of either inducer (Fig. 1c, 1d), meaning the source of the observed growth inhibition likely stems from translational load due to resource limitation^40^. The results show that the degree of metabolic burden is unique to each host and can therefore contribute to the chassis-effect.

Cell densities (OD_600_) were measured simultaneously as fluorescence during the toggle assay to characterize the specific growth rate, lag time and carrying capacity of each host (Fig. 3a and 3b). Both pS4-carrying *Halopseudomonas* spp. revealed the lowest specific growth rates and carrying capacities across induction states, consistent with the WT strains (Fig. 3g). The growth metrics of the three *Pseudomonas* spp. (Fig. 3g) differed appreciably depending on their past induction history, reinforcing the observed host-specific hysteresis effects on growth phenotypes. For instance, *P. deceptionensis* cells toggled from initial OFF to aTc-ON experienced a long lag phase and reduced specific growth rate. But when cells were toggled from Ara-ON to aTc-ON, the prolonged lag phase and growth rate reduction was alleviated, and growth more closely resembled the dynamics of non-induced control cells. A similar hysteresis effect was observed for *P. fluorescens*. It is notable that despite the low PCN, the two *Halopseudomonas* spp. reached the highest SS_aTc-mKate_ values in both toggles, approximately 7-fold higher than the SS_aTc-mKate_ reached by *P. deceptionensis*, the host with the highest PCN. A high PCN therefore does not necessarily result in high SS fluorescent protein levels, but this is also dependent on the promoter or fluorescent protein, as the same trend was not observed for sfGFP expression. The high SS_aTc-mKate_ values obtained by the *Halopseudomonas* spp. could be due to lower rate of cell division and associated dilution of the mKate protein, as the two hosts also exhibit the lowest specific growth rates. This theory is supported by the fact that when *P. deceptionensis* was alleviated of the growth inhibition (increased growth rate), the attained SS_aTc-mKate_ value also decreased by 0.4-fold.

We further considered codon usage bias as an innate host property capable of affecting toggle switch performance, which we parameterized by determining CAI values for the *sfGFP*, *mKate*, *AraC* and *TetR* coding sequences within the toggle switch. Hosts with a codon usage bias more adapted to the synonymous codon usage in the four coding genes (CAI closer to 1) in the toggle switch could lead to more efficient translation of gene products thereby affecting performance, and *vice versa* (CAI closer to 0). *P. deceptionensis* and *P. fluorescens* have the highest CAI values across all four genes (Fig. 3g), *P. putida* on the other hand, has the lowest CAI values across all four genes; a position it shares with *E. coli* for the *AraC* and *TetR* genes only. Previous findings suggest that similar GC content predicts similar codon usage in prokaryotes^41^, yet *E. coli*, with the lowest GC content of 51%, has relatively similar CAI values to *Halopseudomonas* and *Pseudomonas* (GC content 59% - 62%), for all four translated proteins. Our results do not show the expected trend between CAI value and fluorescence output. Still, sequence optimization strategies have been reported to successfully elevate heterologous protein expression^25, 42^, we therefore decided to include the calculated CAI values in downstream analysis.

The summarized physiology metrics can also be abstracted to signature barcodes that collectively represent the hosts physiology profiles. The expression of the toggle switch is inherently coupled to the unique network of metabolic fluxes of each host due to the dependency on host machinery and resources. Given this complexity, it is unlikely that a single variable can reliably predict the performance metrics of our toggle switch. Considering multiple explanatory variables, could provide a more comprehensive prediction of overall toggle switch performance.

### Significant concordance between host physiology and the chassis-effect

The overarching goal of this study was to determine if the chassis-effect is more related to differences in host physiology or phylogenomic relatedness. To formally assess the relationship between species-specific differences in performance, physiology and phylogeny, we employed two multivariate statistical approaches, namely the Mantel test^43, 44^ and Procrustean Superimposition^45–47^ (PS) analysis. Both tests revealed significant concordance only between host physiology and toggle switch performance, indicating that the assorted host physiology metrics serve as a more robust predictor of toggle switch performance than phylogenomic distance-based phylogeny, within our selection of Gammaproteobacteria hosts.

The Mantel test and Procrustean Superimposition (PS) analyses are measures for “goodness-of-fit”, where a high goodness-of-fit indicates the two datasets are correlated. PS analysis acts on ordinated data by centering, scaling, and superimposing two projected ordination configurations to minimize the vector residuals between each item. The Mantel test considers linear correlation between two distance matrices and produces an R statistic that is similar to the Pearson’s Correlation Coefficient; with increasingly similar dissimilarity matrices, the Mantel R statistic will approach 1. PS analysis produces the M^2^ statistic, which is the sum of squared residuals between the best fit of two configurations, such that a M^2^ value closer to 0 indicates a better fit.

To apply the Mantel tests, Euclidean distance matrices were first generated from the respective physiology and performance metric tables (Fig. 4a). Meanwhile the phylogenomic tree was converted to a phylogenomic distance matrix for downstream analysis. The Mantel test revealed a strong significant positive correlation between performance and physiology distance matrices (Fig. 4f, Mantel R = 0.621, p-value = 0.011), indicating that hosts similar in terms of performance are also similar in their overall physiology, and *vice versa*. When testing performance against phylogeny, no significant correlation was observed (Mantel R = 0.160, p = 0.188). The Mantel test summarizes the results in a single metric that only describes the general trend, while PS analysis has the advantage in that the contribution of residuals from each individual item (host) is reported. To investigate the result of the Mantel test further, PS analysis was performed to compare the similarity of the ordination configurations of the hosts.

**Figure 4.**
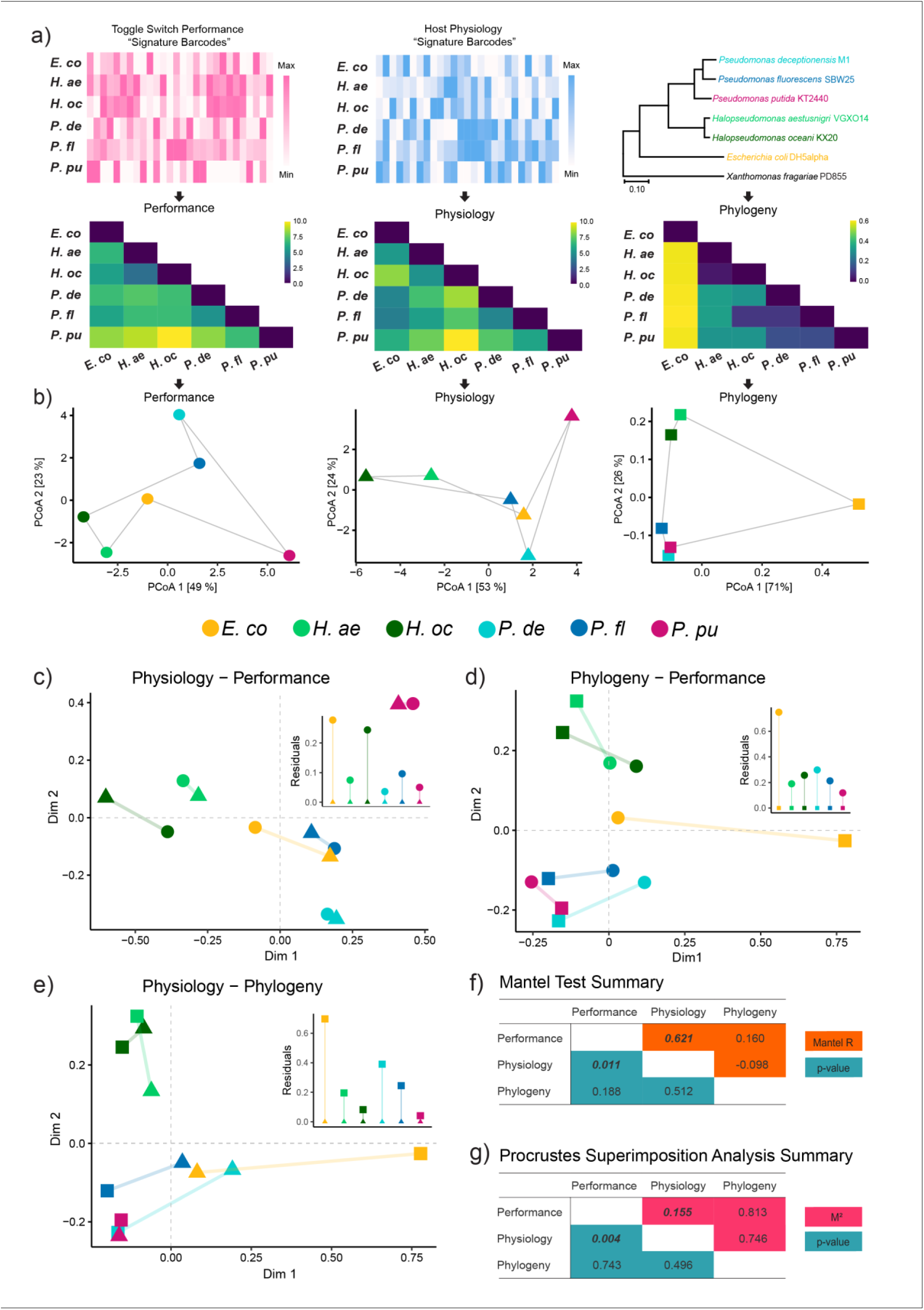
Mantel and Procrustean Superimposition analysis reveal significant concordance between clustering pattern of hosts in terms of physiology and toggle switch performance variation. a) Table of summarized performance (left) and physiology (middle) metrics converted to Euclidean distance metrics (black arrows) with equal weight for each column. Phylogenomic distance matrix was created in MEGAX using the same amino acid MLSA used to infer the phylogenomic tree. b) Principal Coordinate analysis biplots of Performance (circles), Physiology (triangles) and Phylogeny (squares) derived from each respective distance matrix. Hosts are color coded (legend in middle of figure). Solid grey lines were added to help visualize the “configuration” that represents the clustering pattern of hosts for each data type, the lines connect each species in the same arbitrary order. Values in square brackets represent proportion of variation captured by each axis. c) Symmetric pair-wise Procrustean Superimposition analysis of the clustering patterns of each host after scaling, centering and rotating configurations against each other to minimize vector residuals, fitting phylogeny against performance, d) physiology against performance and e) physiology against phylogeny. Hosts are color coded and symbol key follows panel b). Colored lines connecting points represent the vector residuals between respective hosts. Insets shows bar chart of vector residuals. Intersection of dashed gray lines indicates centroid of configurations. Summarized statistics from pair-wise g) Mantel test and h) Procrustean Superimposition analysis. Numbers in bold and italics are significant with p-value < 0.05.

Principal Coordinate Analysis (PCoA) was applied to distance matrices to project the distances in ordination space (Fig. 4b). As expected, ordination of the phylogenomic distance matrix produces a clustering pattern that reflects the branching pattern of the phylogenomic tree (Fig. 4b, right). *Halopseudomonas* spp. clustered across all three types of data, while this is expected in terms of phylogeny, it means they are also similar in terms of their physiology and the toggle switch performance metrics. Pair-wise PS analysis confirmed and strengthened our main result, again reporting significant concordance between the clustering pattern of the hosts in terms of observable variance between host-specific physiology and toggle switch performance (Fig. 4c, M^2^ = 0.155, p = 0.004), while again no significance was found between phylogeny and performance (Fig. 4d, M^2^ = 0.813, p = 0.743). Furthermore, neither test found a significant correlation between physiology and phylogeny (Fig. 4e, f), questioning the commonly assumed predictive power of phylogeny for assessing microbial phenotypes or ecophysiology. Closer inspection of each PS comparison revealed an uneven contribution of residuals by each individual host. In the Physiology – Performance comparison, *E. coli* alone accounts for 36% of the total residual in the data, more than double the theoretical even contribution (16.67%). The second highest contributor of residuals is *H. aestusnigri* with 21% and all other hosts contributing less. This suggests that these two hosts do not follow the significant trend (i.e., physiology underpins performance) as strongly as the other hosts. The reason for this unevenness could be because the underlying mechanisms that drive the chassis-effect in *E. coli* and *H. aestusnigri* were not included in the analysis, although despite this the overall low residual was still deemed significant. When comparing Phylogeny – Performance, *E. coli* is again is responsible for the highest proportion of the residual (41%). This suggests differences in toggle switch performance is trackable along the phylogenomic tree of species part of the *Pseudomonadaceae* family but loses reliability as it approaches *E. coli*.

## DISCUSSION

### Host physiology versus genome relatedness

In this study, we demonstrate within our Gammaproteobacteria framework that the comparative chassis-effects on a genetic toggle switch operated between hosts is better explained by the physiological differences of the host rather than genomic relatedness. In practice, the addition of a new host to our framework could have its toggle switch performance predicted by comparing its quantified physiology against the other characterized hosts. A major caveat of this result is that a given collection of hosts must also share some base-level of genetically compatibility with a given engineered device, therein maintaining some unspecifiable level of dependence upon genetic relatedness. Yet, our results show that the observable chassis-effect cannot be reliably inferred from phylogenomic relationships – i.e., genome relatedness. The results of this study provide a deeper understanding of the biological determinants that manifest a comparable and quantifiable chassis-effect. This study also establishes a framework for testing and predicting how an engineered genetic circuit might perform across closely related hosts and overall bolsters momentum in the field of BHR biodesign. Furthermore, to our knowledge, this study is the first to report the successful genetic engineering of three novel marine hosts, *H. aestusnigri* VGOX14, *H. oceani* KX20 and *P. deceptionensis* M1, with useful innate phenotypes such as polyethylene terephthalate (PET) degradation^48^, psychrotolerance and salinity tolerance^49, 50^.

The diverse range of observed constants across the different chassis was interesting; since both the protein sequences are identical across the data set, changes in kinetic parameters such as the activation constants cannot arise due to differences in binding affinity of the small activating ligands for the transcription factors. Similarly, variations in the Hill coefficient cannot be due to local DNA binding interactions since the promoter sequences are identical. Instead, these chassis-specific effects are more likely due to the expression level of the transcription factor and the bioavailability of the activating ligand, which may also be dependent on transport regulators and efflux. For instance, the markedly higher K_aTc_ values for the *Pseudomonas* spp. could be due to the presence of a native TtgABC efflux pump, known to be more highly expressed in presence of tetracycline in *P. putida* DOT-T1 strain^51, 52^. Indeed, a BLAST search with the *P. putida* DOT-T1 TtgA, TtgB and TtgC genes (NCBI GenBank accession = AF031417.2) against the genomes of our six hosts reveals that all three genes are also present in the *P. putida* and *P. fluorescens* genome, the hosts with the highest K_aTc_. Active export of the inducer leading to lowered intra-cellular concentration could explain the wider induction range of *P. putida* and *P. fluorescens*. This is a clear example of how the presence of a single element can be a major contributor to variation. Furthermore, unpredictable elements such as presence of DNA binding sites on the host genome recognized by the TetR and AraC repressors could lead to sequestration of the transcription factors, leading to differing steady-state concentration of free repressor available to bind to its intended cognate operator^53^. We note that the physiology metrics chosen to contextualize our hosts are by no means exhaustive, and that additional insight could be garnered by measurements that reflects the availability of transcriptional and translational machinery and resources. Examples of such metrics are steady state ribosomal abundance^54^, RNA polymerase abundance and NADH pool^55^. However, the accurate measurement of some of these metrics can be difficult to normalize across hosts^56^.

### Harnessing the chassis-effect

The chassis-effect can manifest in different ways between hosts, from complete circuit inoperability in some hosts to variations in performance between others. The diversity of performance metric profiles exhibited by the six hosts characterized here encourages the notion of viewing chassis as pragmatic parts capable of tuning device parameters. In cases where the tuning of promoter or RBS strengths does not yield the desired functional outcome, exploring chassis design space could be a promising direction. However, we recognize that successfully introducing a device into a novel chassis is often not possible, as exemplified during our design-build-test processes, the pS4 plasmid was not successfully transformed into two initially chosen hosts: *Pseudomonas taeanensis* MS-3 and *Halopseudomonas pachastrellae* CCUG 46540 (NCBI Assembly accession number MS-3 and ASM198937v1, respectively). Whether this was due to transformation methodology, genetic compatibility or the activity of toggle switch being toxic to the cells is unknown, indicating that the “inoperable chassis-effect” is challenging to assess.

### Towards broad-host-range synthetic biology

Armed with automated circuit design software^57^, standardized parts libraries and robust DNA assembly technology, synthetic biologists are now able to reliably manifest their genetic circuit abstractions into physical DNA molecules that carry out the programmed functions with increasing fidelity. We have improved our ability to control for unwanted compositional and context effects^17^ in our circuit designs to a high degree by compartmentalizing modules and components using regulatory elements such as terminators and ribozymes^58^ and rationally avoiding promiscuous genetic elements known to “cross-talk”. However, continued reliance on our limited number of model organisms will surely stagnate the rate of progress of engineering microorganisms. As we domesticate more pragmatic microbes as novel chassis, our ability to control and predict host-context effect must also advance. The emerging field of BHR synthetic biology aims to develop engineering principles that minimize unpredictability arising from host context and our findings contribute to this goal by improving prediction of the chassis-effect; we have here uncovered potential fundamental principles that drives observable differences in host-specific genetic device performance, and thereby lending increased predictive power for porting synthetic circuits across bacterial species and the exploration of chassis design space.

## METHODS

### Strains and Plasmids

WT cells were purchased from the German Collection of Microorganisms and Cell Cultures (DSMZ), except for *Pseudomonas fluorescens* SBW25, which was kindly donated by Rosemarie Wilton from the Argonne National Laboratory (Illinois, USA). Bacterial strains and plasmids used in this study are summarized in Supplementary Table S2, primers used are listed in Supplementary Table S3. All cloning work was performed in *E. coli* DH5α. *E. coli* DH5α cells were made chemically competent and transformed by heat shock following the Inoue method^59^. Species identities were verified by *rpoD* sequencing amplified with primers PsEG30F and PsEG790R as described by Mulet et al. (2019)^60^.

### Media and Culturing Conditions

All bacteria were cultured in Lysogeny-Broth (LB) at 30 °C unless specified. Toggle switch-carrying strains were cultivated with 100 ug/mL kanamycin, while WT counterparts were grown without kanamycin. Overnight cultures were prepared by inoculation with single colonies from streaked plate and cultured with shaking. 96-well plate cultivation was done in black and flat clear-bottom plates (Thermo Fischer, 165305), where 199 µL of media was inoculated with 1 µL of culture and sealed with Breath-Easy film (Sigma-Aldrich, Z380059). OD_600_, sfGFP (485/515, gain = 75 and mKate (585/615, gain = 125) fluorescence was measured continuously up to 48 h using a Synergy H1 plate reader (Agilent Biotek, Serial Number 21031715) with continuous linear shaking (1096 cpm, 1 mm). 1 M Arabinose stocks was prepared dissolving Arabinose (VWR, A11921) in milliQ water and immediately filter sterilized through a 0.2 µM filter and stored at 4 °C. 1 mg/mL aTc stock was prepared by dissolving anhydrotetracycline hydrochloride (VWR, CAYM10009542) powder in 70% ethanol and stored at -20 °C in the dark.

### BASIC Assembly

pS4 toggle switch was assembled in the Biopart Assembly Standard for Idempotent Cloning^29, 30^ (BASIC). For complete protocol, see Storch et al. (2015). DNA parts used in this study are listed in Supplementary Table S4. Briefly, BASIC assembly is a “one-pot” DNA assembly method for joining of DNA parts, following their integration into the BASIC standard. BsaI-HFv2 restriction enzyme and T4 DNA Ligase were purchased from NewEngland Biolabs (R3733 and M0202L, respectively). Mag-Bind TotalPure NGS (Omega Bio-Tek, M1378-01) was used to purify restriction-ligation reactions as per manufacturer’s instructions. BASIC linkers were purchased from Biolegio (BASIC Linkers Pro-Plate Set, BBT-18500). pSEVA231 plasmid was donated by the SEVA repository and was integrated into the BASIC format with primers B_SEVA_F and B_SEVA_R primers (**Error! Reference source not found.**) using the KAPA HiFi HotStart ReadyMix PCR Kit with the following program: initial denaturation at 95 °C for 180 sec, followed by 30 cycles of the following: 98 °C for 20 sec, 58 °C for 15 sec, 72 °C for 150 sec, and final extension step at 72 °C for 150 sec. *AraC*, *TetR*, *sfGFP* and *mKate* genes were ordered from Twist Biosciences with the BASIC *i*P and *i*S sequences integrated at the 5’-end and 3’-end respectively.

### Preparation of Electrocompetent Cells and Electroporation

All incubation steps were done at 30 °C unless specified. 5 mL of overnight culture was diluted by half with fresh LB media and incubated for 1 hour. For *Halopseudomonas oceani* KX20 and *Halopseudomonas aestusnigri* VGXO14, 5 mL of culture was harvested per transformation. For all other species, 1.5 mL was harvested per transformation reaction. Harvesting was done by centrifugation at 4000 RPM at room temperature. Supernatant was discarded and cells resuspended in sucrose electroporation buffer (300 mM sucrose, 1 mM MgCl, pH = 7.2). Washing step was repeated for a total of two washes. Cells were resuspended in 80 µL electroporation buffer and incubated with 50-100 ng of plasmid at room temperature for 15 min. Cells were transferred a 1 mm-gap sterile electroporation cuvette (VWR, 732-1135) and electroporated using an ECM 399 Exponential Decay Wave Electroporation System (BTX, 45-0000) at 1250 V with High Voltage setting (150 resistance and 36 capacitance). 750 µL of LB media was immediately added to the electroporated cells and mixed by pipetting before transferring to 5 mL of LB for recovery by incubation with shaking for 2 h. After recovery, cells were harvested and streaked on kanamycin supplemented LB plates. Colonies would appear after 1 or 2 overnight incubations. Colonies were verified by colony PCR with LMP-F and LMS-R primers.

### Induction Assay

LB media with kanamycin and serial dilutions of inducers Ara and aTc was prepared in separate tubes before aliquoting onto a 96-well plate and inoculated with overnight culture. Normalized steady-state fluorescence intensity was quantified by averaging fluorescence over a time window of 6 - 12 hours at late phase and plotted against induction concentration to yield induction curves. Hill coefficient (n), activation coefficient (K), max steady-state fluorescence output (β) and basal fluorescence output at 0 inducer concentration (y_0_) was estimated by fitting the Hill equation (eq. 1) to induction curves using non-linear least-square regression with the *nls* function from the *stats* base R package with the *port* algorithm.

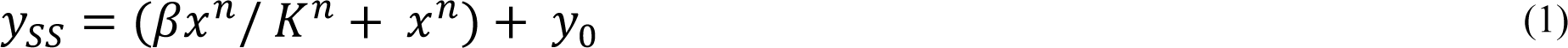

Where y_ss_ is steady-state fluorescence output of sfGFP or mKate at a given inducer concentration and x is Ara (mM) or aTc (ng/mL) inducer concentration.

### Toggle and Growth Assay

Based on the induction assay, arbitrary inducer concentration of 20 mM Ara and 10 ng/mL aTc was chosen to induce all species. Overnight cultures grown in absence of inducer were used to inoculate media in 96-well plate supplemented with Ara and aTc. Media without inducer was also inoculated as control. OD_600_, sfGFP and mKate fluorescence was continuously measured. To toggle, cells were harvested by centrifugation at 4000 RPM for 20 minutes at room temperature and supernatant discarded before resuspending in equal volume LB media, this washing step was repeated for a total of two washes. 1 µL of washed resuspended cells were inoculated to 199 µL fresh media supplemented with the opposite respective inducer. Metrics Δµ_NI_, Δµ_Ara_ and Δµ_aTc_ were calculated with equation 2, 3 and 4 respectively.

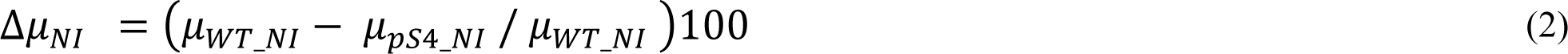

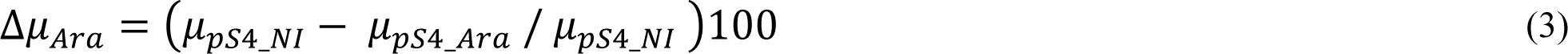

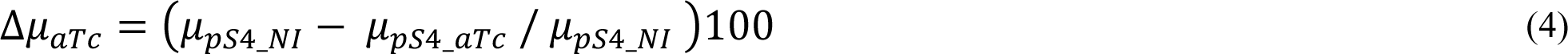

Where µ_WT_NI_ is the specific growth rate of the WT strain without inducer and µ_pS4_NI_, µ_pS4_Ara_ and µ_pS4_aTc_ are the specific growth rates of the pS4-carrying counterpart strain in absence of inducer, in presence of 20 mM Ara and in presence of 10 ng/mL aTc, respectively. Maximum slopes of OD_600_ and fluorescence curves were estimated based on a rolling regression method using the *all_easylinear* function from the *growthrates* (v.0.8.4, https://CRAN.R-project.org/package=growthrates) R package. The Lag times and curve plateaus of OD_600_ and fluorescence curves were determined using the *all_growthmodels* function, fitting the Gompertz growth model^61^ with additional lag (lambda) parameter.

### Plasmid Copy Number Determination

PCN per chromosome was determined through real-time PCR method based on Škulj, et al. (2008)^36^ using the single-copy *rpoD* gene and the *bla* gene in pS4 as amplification targets. Primers for qPCR were designed with BatchPrimer3^62^ with the following criteria: product size = 100 - 125 bp, primer size = 20 - 24 bp, primer Tm = 58 - 62 °C, primer GC % = 45-55 %. Cells were harvested at stationary phase (18 – 24 h) by centrifugation at 4500 rpm, 4 °C for 10 min and resuspended to OD_600_ = 1 in cold PBS (1x, pH = 7.3). Up to 1 mL of resuspended culture was incubated at 95°C for 20 min to lyse the cells and immediately stored at -20 °C until use as template for real-time PCR. Simultaneously, resuspended culture was serially diluted to 10^-9^ and 100 μl of 10^-9^, 10^-8^, 10^-7^ and 10^-6^ dilutions was streaked on solid LB agar medium with kanamycin and incubated overnight at 30 °C for viable cell count. Lysed template samples were diluted to dynamic range of 10^2^ – 10^5^ bacteria per well. Real-Time PCR reactions were performed in 20 µL mixtures containing 10 µL KAPA SYBR FAST qPCR Master Mix (2X) (Sigma-Aldrich, KK4605), 200 nM final concentration of species-specific *rpoD* forward and reverse primer and up to 3 µL of sample template. Separate reactions were done for chromosomal and plasmid amplicons, each in four technical replicates. Reaction solutions were aliquoted onto a fast optical 0.1 µL 96-well plate (Applied Biosystems, 4346906). Real-time PCR was performed using a 7500 Fast Real-Time PCR System (Applied Biosystems, 4351107) with following cycling conditions: 1 min at 95°C followed by 40 cycles of 3 s at 95°C and 30 s at 60°C. A melt-curve analysis step was added with program: 15 s at 95°C, 1 min at 60°C followed by a 1% gradual increase in temperature to 95°C and 15 s at 60°C. Cycle threshold (Ct) values were determined after automatic adjustment of the baseline and fluorescence thresholds in 7500 Software v2.3 (Applied Biosystems). Relative standard curves were constructed in R by plotting log value of the number of colony-forming units against Ct values. Slopes were determined from curves with three to five dilutions and on condition that r^2^ ≥ 0.99. Amplification efficiency (E) for plasmid and *rpoD* was calculated, which was used in turn to determine PCN using equation (6), which takes into consideration different amplification efficiencies. PCN was calculated with the lowest Ct value and mean calculated from three biological replicates.

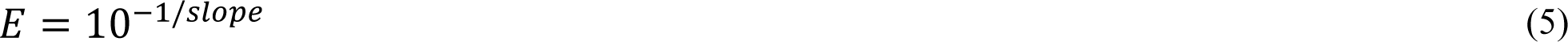

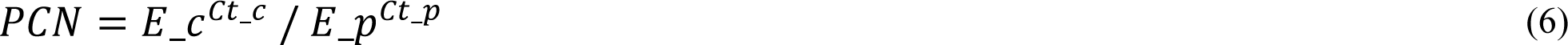

Where E_c is amplification efficiency for *rpoD* gene (chromosome) and E_p is amplification efficiency for *bla* gene (plasmid).

### Phylogenomic and Phylogenetic Tree Building

Genomes were downloaded from the NCBI Assembly database, see Supplementary Table S2 for genome accession numbers. GToTree^33^ (v.1.7.07, https://github.com/AstrobioMike/GToTree) was run with default settings using the Gammaproteobacteria HMM set for 172 orthologous single-copy genes to create an amino-acid MLSA. Tree inference and distance matrix construction was done in MEGAX^63^ using Neighbor-Joining method with Jones-Taylor-Thornton matrix-based model^64^ with uniform rates.

Ambiguous positions were removed for each sequence pair (pairwise deletion option) and standard error estimates were obtained with bootstrap procedure of 1000 replicates.

### Codon Adaption Index

The synonymous codon usage bias of genes encoding ribosomal proteins and ribosomal RNA genes was used as reference for each host, which are genes assumed to be highly expressed and thus under selection pressure to using with codons adapted to the codon usage bias of the organism^35, 65^. Sequences of ribosomal proteins, 16S and 23S rRNA were retrieved using *anvi-get-sequences-for-hmm-hits* function with the *Bacteria_71* HMM set in the anvi’o^66^ (v.7.1, https://github.com/merenlab/anvio) environment. The CAI for the *sfGFP*, *mKate*, *AraC* and *TetR* genes were calculated using the standard genetic code with the CAI python package^67^ (v.1.0.2, https://github.com/Benjamin-Lee/CodonAdaptationIndex). CAI is the geometric mean of the relative adaptiveness of each codon in a given sequence with the following implementation in the CAI python program:

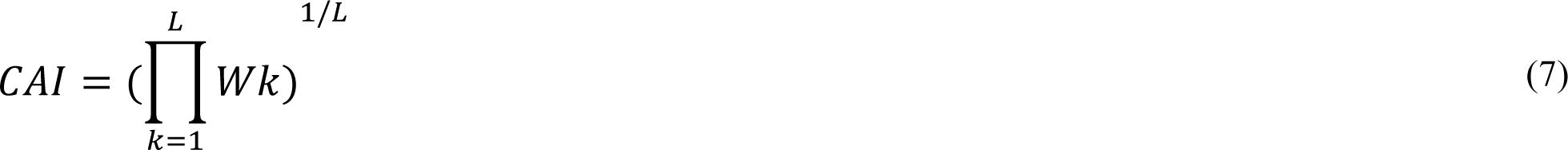

*L* is the number of codons, and *Wk* relative adaptiveness of codon *k* and is calculated as follows:

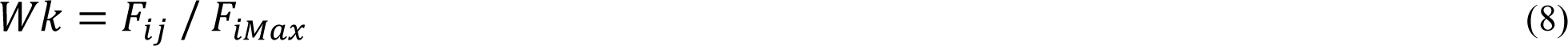

F_ij_ is the frequency of codon *j* for coding for amino acid *i* in a given sequence. F_iMax_ is the frequency of the most optimal codon for amino acid *i* in the given sequence, with the most optimal codon being the one with the highest frequency in the reference set of genes.

### Statistical Analyses

Generation of Euclidean distance matrices (equal weights), PCoA (equal weights), Mantel test and Procrustean Superimposition analysis and were all done using the Vegan^68^ (v.2.6-4, https://CRAN.R-project.org/package=vegan) package in R. Procrustean M^2^ statistic (*scale* and *symmetric* true) and Mantel R statistic (*pearson* method) was tested for significance by a permutation approach (n = 719, max number of iterations). Briefly, observations in one matrix are randomly reordered while maintaining the covariance structure within the matrix and test statistic is calculated and recorded enough times to obtain a sizeable null distribution. A p-value for each statistic is then calculated, representing the probability of obtaining a statistic with value equal to or more “extreme” of the experimental value.

## DATA AND CODE AVAILIBILITY

All experimental data files, raw data files, R MarkDown scripts used for analysis and producing graphs and plots are publicly available online on the Open Science Framework as part of the project name *Chan.Interspecies.Toggle.Switch* (https://osf.io/jnkrx/). Genome, plasmid, and bacterial strain accession numbers can be found in supplementary material.

## ACKNOWLEDGEMENTS

The authors would like to thank Rosemarie Wilton from the Argonne National Laboratory for their donation of the *Pseudomonas fluorescence* SBW25 strain. We also thank the Standard European Vector Architecture (SEVA) for their donation of backbone plasmids. This work was supported by strategic funding from UiT – The Arctic University of Norway under the project ABSORB (Arctic Carbon Storage from Biomes).

## ETHICS DECLERATION

The authors have no conflicts of interest to declare.

## SUPPLEMENTARY MATERIAL

**Supplementary Table S1.** Single copy gene hits of the 172 Gammaproteobacteria Hidden Markov Models in the GToTree Program and report statistics for each species used in this study. The number of hits for each gene is indicated.

|  | Host | *Pseudomonas fluorescens* | *Pseudomonas putida* | *Pseudomonas deceptionensis* | *Halopseudomonas aestusnigri* | *Escherichia coli* | *Halopseudomonas oceani* | *Xanthomonas fragariae* |
| --- | --- | --- | --- | --- | --- | --- | --- | --- |
|  | taxid | #### | #### | #### | #### | 562 | #### | #### |
|  | num_SCG_hits | 166 | 171 | 172 | 172 | 172 | 171 | 167 |
|  | uniq_SCG_hits | 159 | 167 | 171 | 170 | 172 | 167 | 167 |
|  | perc_comp | 97 | 99 | 100 | 100 | 100 | 99 | 97 |
|  | perc_redund | 4.7 | 2.3 | 0.6 | 1.2 | 0 | 2.9 | 0 |
|  | num_SCG_hits_after_len_filt | 159 | 166 | 170 | 170 | 166 | 167 | 159 |
|  | in_final_tree | Yes | Yes | Yes | Yes | Yes | Yes | Yes |
| PFAM Accession Number: | NCBI species | *Pseudomonas fluorescens* | *Pseudomonas putida* | *Pseudomonas deceptionensis* | *Halopseudomonas aestusnigri* | *Escherichia coli* | *Halopseudomonas oceani* | *Xanthomonas fragariae* |
| PF01812.20 | 5-FTHF_cyc-lig | 1 | 1 | 1 | 1 | 1 | 1 | 1 |
| PF00022.19 | Actin | 1 | 1 | 1 | 1 | 1 | 1 | 1 |
| PF00709.21 | Adenylsucc_synt | 1 | 1 | 1 | 1 | 1 | 1 | 1 |
| PF00406.22 | ADK | 1 | 1 | 1 | 1 | 1 | 1 | 1 |
| PF01808.18 | AICARFT_IMPC Has | 1 | 1 | 1 | 1 | 1 | 1 | 1 |
| PF00731.20 | AIRC | 1 | 1 | 1 | 1 | 1 | 1 | 1 |
| PF00490.21 | ALAD | 2 | 2 | 1 | 1 | 1 | 1 | 1 |
| PF03702.14 | AnmK | 1 | 1 | 1 | 1 | 1 | 1 | 1 |
| PF00231.19 | ATP-synt | 1 | 1 | 1 | 1 | 1 | 1 | 1 |
| PF00119.20 | ATP-synt_A | 1 | 1 | 1 | 1 | 1 | 1 | 1 |
| PF00430.18 | ATP-synt_B | 1 | 1 | 1 | 1 | 1 | 1 | 1 |
| PF00137.21 | ATP-synt_C | 1 | 1 | 1 | 1 | 1 | 1 | 1 |
| PF02823.16 | ATP-synt_DE_N | 1 | 1 | 1 | 1 | 1 | 1 | 1 |
| PF03899.15 | ATP-synt_I | 1 | 1 | 1 | 1 | 1 | 1 | 0 |
| PF03668.15 | ATP_bind_2 | 1 | 1 | 1 | 1 | 1 | 1 | 1 |
| PF04380.13 | BMFP | 1 | 1 | 1 | 1 | 1 | 1 | 1 |
| PF01264.21 | Chorismate_synt | 1 | 1 | 1 | 1 | 1 | 1 | 1 |
| PF02674.16 | Colicin_V | 1 | 1 | 1 | 1 | 1 | 1 | 1 |
| PF01218.18 | Coprogen_oxidas | 1 | 1 | 1 | 1 | 1 | 1 | 1 |
| PF00166.21 | Cpn10 | 1 | 1 | 1 | 1 | 1 | 1 | 1 |
| PF00118.24 | Cpn60_TCP1 | 1 | 1 | 1 | 1 | 1 | 1 | 1 |
| PF05173.14 | DapB_C | 1 | 1 | 1 | 1 | 1 | 1 | 1 |
| PF01678.19 | DAP_epimerase | 3 | 2 | 1 | 1 | 1 | 1 | 1 |
| PF01761.20 | DHQ_synthase | 1 | 1 | 1 | 1 | 1 | 1 | 1 |
| PF04977.15 | DivIC | 1 | 1 | 1 | 1 | 1 | 1 | 1 |
| PF00885.19 | DMRL_synthase | 2 | 1 | 2 | 2 | 1 | 2 | 1 |
| PF04364.13 | DNA_pol3_chi | 1 | 1 | 1 | 1 | 1 | 1 | 1 |
| PF06144.13 | DNA_pol3_delta | 1 | 1 | 1 | 1 | 1 | 1 | 1 |
| PF02622.15 | DUF179 | 1 | 1 | 1 | 1 | 1 | 1 | 1 |
| PF04241.15 | DUF423 | 1 | 1 | 1 | 1 | 1 | 1 | 1 |
| PF04356.12 | DUF489 | 1 | 1 | 1 | 1 | 1 | 1 | 1 |
| PF04359.14 | DUF493 | 1 | 1 | 1 | 1 | 1 | 1 | 1 |
| PF04751.14 | DUF615 | 1 | 1 | 1 | 1 | 1 | 1 | 1 |
| PF00889.19 | EF_TS | 1 | 1 | 1 | 1 | 1 | 1 | 1 |
| PF01176.19 | eIF-1a | 1 | 1 | 1 | 1 | 1 | 1 | 1 |
| PF00113.22 | Enolase_C | 2 | 1 | 1 | 1 | 1 | 1 | 1 |
| PF02601.15 | Exonuc_VII_L | 1 | 1 | 1 | 1 | 1 | 1 | 1 |
| PF02609.16 | Exonuc_VII_S | 1 | 1 | 1 | 1 | 1 | 1 | 1 |
| PF00762.19 | Ferrochelatase | 1 | 1 | 1 | 1 | 1 | 1 | 1 |
| PF04999.13 | FtsL | 1 | 1 | 1 | 1 | 1 | 1 | 1 |
| PF04551.14 | GcpE | 1 | 1 | 1 | 1 | 1 | 1 | 1 |
| PF01025.19 | GrpE | 1 | 1 | 1 | 1 | 1 | 1 | 1 |
| PF01725.16 | Ham1p_like | 1 | 1 | 1 | 1 | 1 | 1 | 1 |
| PF02602.15 | HEM4 | 1 | 1 | 1 | 2 | 1 | 2 | 1 |
| PF17209.3 | Hfq | 1 | 1 | 1 | 1 | 1 | 1 | 1 |
| PF01634.18 | HisG | 1 | 1 | 1 | 1 | 1 | 1 | 1 |
| PF00815.20 | Histidinol_dh | 2 | 1 | 1 | 1 | 1 | 1 | 1 |
| PF01430.19 | HSP33 | 1 | 1 | 1 | 1 | 1 | 1 | 1 |
| PF00475.18 | IGPD | 1 | 1 | 1 | 1 | 1 | 1 | 1 |
| PF01715.17 | IPPT | 1 | 1 | 1 | 1 | 1 | 1 | 1 |
| PF01745.16 | IPT | 1 | 1 | 1 | 1 | 1 | 0 | 1 |
| PF04362.14 | Iron_traffic | 1 | 1 | 1 | 1 | 1 | 1 | 1 |
| PF06305.11 | LapA_dom | 0 | 1 | 1 | 1 | 1 | 1 | 1 |
| PF03588.14 | Leu_Phe_trans | 1 | 1 | 1 | 1 | 1 | 1 | 1 |
| PF03550.14 | LolB | 1 | 1 | 1 | 1 | 1 | 1 | 1 |
| PF02684.15 | LpxB | 1 | 1 | 1 | 1 | 1 | 1 | 1 |
| PF03331.13 | LpxC | 1 | 1 | 1 | 1 | 1 | 1 | 1 |
| PF02606.14 | LpxK | 1 | 1 | 1 | 1 | 1 | 1 | 1 |
| PF02401.18 | LYTB | 1 | 1 | 1 | 1 | 1 | 1 | 1 |
| PF03776.14 | MinE | 1 | 1 | 1 | 1 | 1 | 1 | 1 |
| PF05494.12 | MlaC | 1 | 1 | 1 | 1 | 1 | 1 | 1 |
| PF12631.7 | MnmE_helical | 1 | 1 | 1 | 1 | 1 | 1 | 1 |
| PF04085.14 | MreC | 1 | 1 | 1 | 1 | 1 | 1 | 1 |
| PF04093.12 | MreD | 1 | 1 | 1 | 1 | 1 | 1 | 1 |
| PF00334.19 | NDK | 1 | 1 | 1 | 1 | 1 | 1 | 1 |
| PF03938.14 | OmpH | 1 | 1 | 1 | 1 | 1 | 1 | 0 |
| PF00213.18 | OSCP | 1 | 1 | 1 | 1 | 1 | 1 | 1 |
| PF02569.15 | Pantoate_ligase | 1 | 1 | 1 | 1 | 1 | 1 | 1 |
| PF02153.17 | PDH | 2 | 1 | 1 | 1 | 1 | 1 | 1 |
| PF00800.18 | PDT | 1 | 1 | 1 | 1 | 1 | 1 | 1 |
| PF03740.13 | PdxJ | 1 | 1 | 1 | 1 | 1 | 1 | 1 |
| PF00311.17 | PEPcase | 1 | 1 | 1 | 1 | 1 | 1 | 1 |
| PF01252.18 | Peptidase_A8 | 1 | 1 | 1 | 1 | 1 | 3 | 1 |
| PF01195.19 | Pept_tRNA_hydro | 1 | 1 | 1 | 1 | 1 | 1 | 1 |
| PF00342.19 | PGI | 1 | 2 | 1 | 1 | 1 | 1 | 1 |
| PF00162.19 | PGK | 1 | 1 | 1 | 1 | 1 | 1 | 1 |
| PF03831.14 | PhnA | 1 | 1 | 1 | 1 | 1 | 1 | 1 |
| PF02233.16 | PNTB | 1 | 1 | 1 | 1 | 1 | 1 | 0 |
| PF01379.20 | Porphobil_deam | 1 | 1 | 1 | 1 | 1 | 1 | 1 |
| PF00697.22 | PRAI | 1 | 1 | 1 | 1 | 1 | 1 | 1 |
| PF01255.19 | Prenyltransf | 1 | 1 | 1 | 1 | 1 | 1 | 1 |
| PF01416.20 | PseudoU_synth_1 | 1 | 1 | 1 | 1 | 1 | 1 | 1 |
| PF02666.15 | PS_Dcarboxylase | 1 | 1 | 1 | 1 | 1 | 1 | 1 |
| PF02033.18 | RBFA | 1 | 1 | 1 | 1 | 1 | 1 | 1 |
| PF02565.15 | RecO_C | 1 | 1 | 1 | 1 | 1 | 1 | 1 |
| PF02631.16 | RecX | 1 | 1 | 1 | 1 | 1 | 1 | 1 |
| PF00825.18 | Ribonuclease_P | 0 | 1 | 1 | 1 | 1 | 1 | 1 |
| PF00687.21 | Ribosomal_L1 | 1 | 1 | 1 | 1 | 1 | 1 | 1 |
| PF00572.18 | Ribosomal_L13 | 1 | 1 | 1 | 1 | 1 | 1 | 1 |
| PF00238.19 | Ribosomal_L14 | 1 | 1 | 1 | 1 | 1 | 1 | 1 |
| PF00252.18 | Ribosomal_L16 | 1 | 1 | 1 | 1 | 1 | 1 | 1 |
| PF01196.19 | Ribosomal_L17 | 1 | 1 | 1 | 1 | 1 | 1 | 1 |
| PF00861.22 | Ribosomal_L18p | 1 | 1 | 1 | 1 | 1 | 1 | 1 |
| PF01245.20 | Ribosomal_L19 | 1 | 1 | 1 | 1 | 1 | 1 | 1 |
| PF00453.18 | Ribosomal_L20 | 1 | 1 | 1 | 1 | 1 | 1 | 1 |
| PF00829.21 | Ribosomal_L21p | 1 | 1 | 1 | 1 | 1 | 1 | 1 |
| PF00237.19 | Ribosomal_L22 | 1 | 1 | 1 | 1 | 1 | 1 | 1 |
| PF00276.20 | Ribosomal_L23 | 1 | 1 | 1 | 1 | 1 | 1 | 1 |
| PF17136.4 | ribosomal_L24 | 1 | 1 | 1 | 1 | 1 | 1 | 1 |
| PF01016.19 | Ribosomal_L27 | 1 | 1 | 1 | 1 | 1 | 1 | 1 |
| PF00828.19 | Ribosomal_L27A | 1 | 1 | 1 | 1 | 1 | 1 | 1 |
| PF00830.19 | Ribosomal_L28 | 1 | 1 | 1 | 1 | 1 | 1 | 1 |
| PF00831.23 | Ribosomal_L29 | 1 | 1 | 1 | 1 | 1 | 1 | 1 |
| PF00297.22 | Ribosomal_L3 | 1 | 1 | 1 | 1 | 1 | 1 | 1 |
| PF01783.23 | Ribosomal_L32p | 1 | 1 | 1 | 1 | 1 | 1 | 1 |
| PF00471.20 | Ribosomal_L33 | 1 | 1 | 1 | 1 | 1 | 1 | 1 |
| PF00468.17 | Ribosomal_L34 | 0 | 1 | 1 | 1 | 1 | 1 | 1 |
| PF01632.19 | Ribosomal_L35p | 1 | 1 | 1 | 1 | 1 | 1 | 1 |
| PF00573.22 | Ribosomal_L4 | 1 | 1 | 1 | 1 | 1 | 1 | 1 |
| PF00347.23 | Ribosomal_L6 | 1 | 1 | 1 | 1 | 1 | 1 | 1 |
| PF03948.14 | Ribosomal_L9_C | 1 | 1 | 1 | 1 | 1 | 1 | 1 |
| PF00338.22 | Ribosomal_S10 | 1 | 1 | 1 | 1 | 1 | 1 | 1 |
| PF00411.19 | Ribosomal_S11 | 1 | 1 | 1 | 1 | 1 | 1 | 1 |
| PF00416.22 | Ribosomal_S13 | 1 | 1 | 1 | 1 | 1 | 1 | 1 |
| PF00253.21 | Ribosomal_S14 | 1 | 1 | 1 | 1 | 1 | 1 | 1 |
| PF00312.22 | Ribosomal_S15 | 1 | 1 | 1 | 1 | 1 | 1 | 1 |
| PF00886.19 | Ribosomal_S16 | 1 | 1 | 1 | 1 | 1 | 1 | 1 |
| PF00366.20 | Ribosomal_S17 | 1 | 1 | 1 | 1 | 1 | 1 | 1 |
| PF00203.21 | Ribosomal_S19 | 1 | 1 | 1 | 1 | 1 | 1 | 1 |
| PF00318.20 | Ribosomal_S2 | 1 | 1 | 1 | 1 | 1 | 1 | 1 |
| PF01649.18 | Ribosomal_S20p | 1 | 1 | 1 | 1 | 1 | 1 | 1 |
| PF01165.20 | Ribosomal_S21 | 1 | 1 | 1 | 1 | 1 | 1 | 1 |
| PF01250.17 | Ribosomal_S6 | 1 | 1 | 1 | 1 | 1 | 1 | 1 |
| PF00177.21 | Ribosomal_S7 | 1 | 1 | 1 | 1 | 1 | 1 | 1 |
| PF00410.19 | Ribosomal_S8 | 1 | 1 | 1 | 1 | 1 | 1 | 1 |
| PF00380.19 | Ribosomal_S9 | 1 | 1 | 1 | 1 | 1 | 1 | 1 |
| PF00164.25 | Ribosomal_S12_S23 | 1 | 1 | 1 | 1 | 1 | 1 | 1 |
| PF06026.14 | Rib_5-P_isom_A | 1 | 1 | 1 | 1 | 1 | 1 | 1 |
| PF02646.16 | RmuC | 1 | 1 | 1 | 1 | 1 | 1 | 1 |
| PF01351.18 | RNase_HII | 1 | 1 | 1 | 1 | 1 | 1 | 1 |
| PF01193.24 | RNA_pol_L | 1 | 1 | 1 | 1 | 1 | 1 | 1 |
| PF01192.22 | RNA_pol_Rpb6 | 1 | 1 | 1 | 1 | 1 | 1 | 1 |
| PF01765.19 | RRF | 1 | 1 | 1 | 1 | 1 | 1 | 1 |
| PF02410.15 | RsfS | 1 | 1 | 1 | 1 | 1 | 1 | 1 |
| PF02075.17 | RuvC | 1 | 1 | 1 | 1 | 1 | 1 | 1 |
| PF03652.15 | RuvX | 1 | 1 | 1 | 1 | 1 | 1 | 1 |
| PF01259.18 | SAICAR_synt | 1 | 1 | 1 | 1 | 1 | 1 | 1 |
| PF04445.13 | SAM_MT | 1 | 1 | 1 | 1 | 1 | 1 | 0 |
| PF03937.16 | Sdh5 | 1 | 1 | 1 | 1 | 1 | 1 | 1 |
| PF02556.14 | SecB | 1 | 1 | 1 | 1 | 1 | 1 | 1 |
| PF00584.20 | SecE | 0 | 1 | 1 | 1 | 1 | 1 | 1 |
| PF03840.14 | SecG | 0 | 1 | 1 | 1 | 1 | 1 | 1 |
| PF00344.20 | SecY | 1 | 1 | 1 | 1 | 1 | 1 | 1 |
| PF04102.12 | SlyX | 1 | 1 | 1 | 1 | 1 | 1 | 1 |
| PF01668.18 | SmpB | 1 | 1 | 1 | 1 | 1 | 1 | 1 |
| PF02590.17 | SPOUT_MTase | 1 | 1 | 1 | 1 | 1 | 1 | 1 |
| PF04386.13 | SspB | 1 | 1 | 1 | 1 | 1 | 1 | 1 |
| PF00902.18 | TatC | 1 | 2 | 1 | 1 | 1 | 1 | 1 |
| PF01702.18 | TGT | 1 | 1 | 1 | 1 | 1 | 1 | 1 |
| PF00303.19 | Thymidylat_synt | 1 | 1 | 1 | 1 | 1 | 1 | 1 |
| PF03966.16 | Trm112p | 1 | 1 | 1 | 1 | 1 | 1 | 1 |
| PF00750.19 | tRNA-synt_1d | 1 | 1 | 1 | 1 | 1 | 1 | 1 |
| PF01411.19 | tRNA-synt_2c | 2 | 1 | 1 | 1 | 1 | 1 | 1 |
| PF02091.15 | tRNA-synt_2e | 1 | 1 | 1 | 1 | 1 | 1 | 1 |
| PF01746.21 | tRNA_m1G_MT | 1 | 1 | 1 | 1 | 1 | 1 | 1 |
| PF02092.17 | tRNA_synt_2f | 1 | 1 | 1 | 1 | 1 | 1 | 1 |
| PF02580.16 | Tyr_Deacylase | 1 | 1 | 1 | 1 | 1 | 1 | 1 |
| PF03658.14 | Ub-RnfH | 1 | 0 | 1 | 1 | 1 | 1 | 1 |
| PF02130.17 | UPF0054 | 1 | 1 | 1 | 1 | 1 | 1 | 1 |
| PF02021.17 | UPF0102 | 1 | 1 | 1 | 1 | 1 | 1 | 1 |
| PF17775.1 | UPF0225 | 1 | 1 | 1 | 1 | 1 | 1 | 1 |
| PF14681.6 | UPRTase | 1 | 1 | 1 | 1 | 1 | 2 | 1 |
| PF01208.17 | URO-D | 1 | 1 | 1 | 1 | 1 | 1 | 1 |
| PF02699.15 | YajC | 1 | 1 | 1 | 1 | 1 | 1 | 1 |
| PF02575.16 | YbaB_DNA_bd | 1 | 1 | 1 | 1 | 1 | 1 | 1 |
| PF02620.17 | YceD | 1 | 1 | 1 | 1 | 1 | 1 | 1 |
| PF02618.16 | YceG | 1 | 1 | 1 | 1 | 1 | 1 | 1 |
| PF05166.13 | YcgL | 1 | 1 | 1 | 1 | 1 | 1 | 1 |
| PF02542.16 | YgbB | 1 | 1 | 1 | 1 | 1 | 1 | 1 |
| PF02325.17 | YGGT | 1 | 1 | 1 | 1 | 1 | 1 | 0 |
| PF03755.13 | YicC_N | 1 | 1 | 1 | 1 | 1 | 1 | 1 |
| PF01809.18 | YidD | 0 | 1 | 1 | 1 | 1 | 1 | 1 |

**Supplementary Table S2.** Metadata on strains used in this study. DSM strain number and NCBI Assembly accession numbers provided are the latest at the time of the publication of this study.

| Species | Genotype | Culture Collection* | NCBI Accession | Ref. |
| --- | --- | --- | --- | --- |
| *Halopseudomonas aestusnigri* VGXO14 | WT | DSMZ (103065) | ASM219798v1 | 1,2 |
| *Pseudomonas deceptionensis* M1 | WT | DSMZ (26521) | G8684 | 3,4 |
| *Pseudomonas fluorescens* SBW25 | WT | Donated** | MPBAS00001 | 5 |
| *Halopseudomonas oceani* KX20 | WT | DSMZ (100277) | ASM290316v1 | 6,7 |
| *Pseudomonas putida* KT2440 | WT | DSMZ (6125) | ASM756v2 | 8 |
| *Escherichia coli* DH5α | WT | DSMZ (6897) | ASM289947v1 | 9 |
| *Xanthomonas fragariae* PD855 | WT | Na | PD885.1 | 10 |
| *Halopseudomonas aestusnigri* VGXO14 | pS4 | This study | This study | This study |
| *Pseudomonas deceptionensis* M1 | pS4 | This study | This study | This study |
| *Pseudomonas fluorescens* SBW25 | pS4 | This study | This study | This study |
| *Halopseudomonas oceani* KX20 | pS4 | This study | This study | This study |
| *Pseudomonas putida* KT2440 | pS4 | This study | This study | This study |
| *Escherichia coli* DH5α | pS4 | This study | This study | This study |

\*DSMZ = German Collection of Microorganisms and Cell Cultures. Numbers in brackets are DSMZ
Culture Collection accession numbers.
\*\*Donation by Rosemarie Wilton, Argonne National Laboratory

**Supplementary Table S3.**
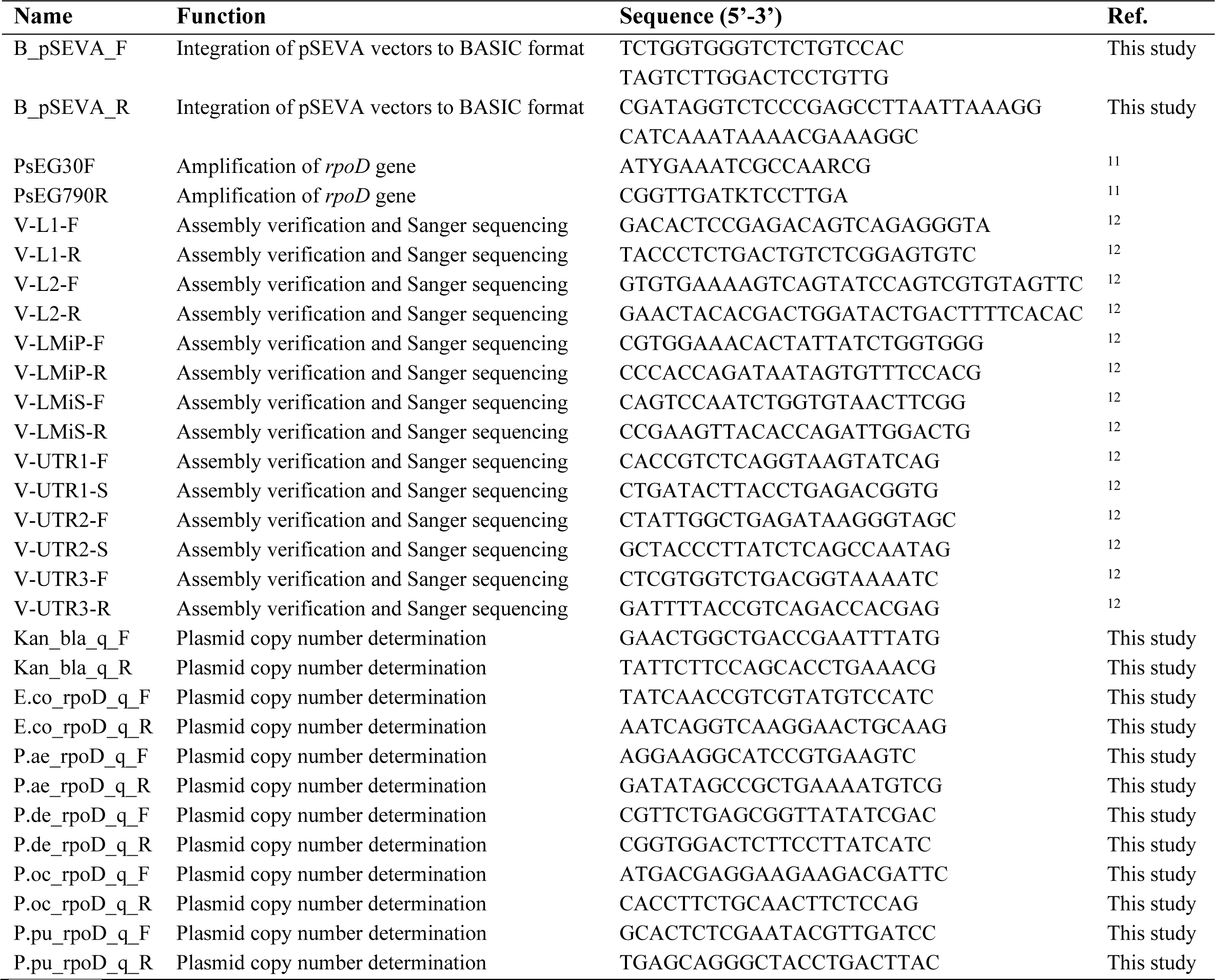
Primers used in this study.

**Supplementary Table S4.** DNA parts and BASIC linkers used in BASIC assembly of pS4 plasmid this study. Accession numbers provided are the latest at the time of the publication of this study.

| Gene/Part | Function | NCBI Accession | Cloning | Ref. |
| --- | --- | --- | --- | --- |
| sfGFP | Fluorescent reporter gene | MG323875 | Synthesized from BioTwist | 13 |
| mKate | Fluorescent reporter gene | MN623117 | Synthesized from BioTwist | 13 |
| AraC | Gene encoding L-Ara inducible repressor | MH101733 | Synthesized from BioTwist | 14 |
| TetR | Gene encoding aTc inducible repressor | MH101732 | Synthesized from BioTwist | 14 |
| B_23 | BASIC integrated pSEVA231 backbone with pBBR1 *ori* and Kan<sup>R</sup> marker | JX560328 | Cloned with B_pSEVA_F and B_pSEVA_R primers | 15 |
| P27_PF1 | Insulated P<sub>Tet</sub> promoter with upstream terminator and riboJ autocatalytic ribozyme | NA | See ref | 16 |
| P32_PF1 | Insulated P<sub>BAD</sub> promoter with upstream terminator and riboJ autocatalytic ribozyme | NA | See ref | 16 |
| BASIC Linkers | BASIC assembly of parts and adding ribosome binding sites | NA | Purchased from BIOLEGIO (BBT-18500) | 12 |

